# Glycosaminoglycan binding motif at S1/S2 proteolytic cleavage site on spike glycoprotein may facilitate novel coronavirus (SARS-CoV-2) host cell entry

**DOI:** 10.1101/2020.04.14.041459

**Authors:** So Young Kim, Weihua Jin, Amika Sood, David W. Montgomery, Oliver C. Grant, Mark M. Fuster, Li Fu, Jonathan S. Dordick, Robert J. Woods, Fuming Zhang, Robert J. Linhardt

## Abstract

Severe acute respiratory syndrome-related coronavirus 2 (SARS-CoV-2) has resulted in a pandemic and continues to spread around the globe at an unprecedented rate. To date, no effective therapeutic is available to fight its associated disease, COVID-19. Our discovery of a novel insertion of glycosaminoglycan (GAG)-binding motif at S1/S2 proteolytic cleavage site (681-686 (PRRARS)) and two other GAG-binding-like motifs within SARS-CoV-2 spike glycoprotein (SGP) led us to hypothesize that host cell surface GAGs might be involved in host cell entry of SARS-CoV-2. Using a surface plasmon resonance direct binding assay, we found that both monomeric and trimeric SARS-CoV-2 spike more tightly bind to immobilized heparin (K_D_ = 40 pM and 73 pM, respectively) than the SARS-CoV and MERS-CoV SGPs (500 nM and 1 nM, respectively). In competitive binding studies, the IC_50_ of heparin, tri-sulfated non-anticoagulant heparan sulfate, and non-anticoagulant low molecular weight heparin against SARS-CoV-2 SGP binding to immobilized heparin were 0.056 μM, 0.12 μM, and 26.4 μM, respectively. Finally, unbiased computational ligand docking indicates that heparan sulfate interacts with the GAG-binding motif at the S1/S2 site on each monomer interface in the trimeric SARS-CoV-2 SGP, and at another site (453-459 (YRLFRKS)) when the receptor-binding domain is in an open conformation. Our study augments our knowledge in SARS-CoV-2 pathogenesis and advances carbohydrate-based COVID-19 therapeutic development.

## Introduction

In March 2020, the World Health Organization declared severe acute respiratory syndrome-related coronavirus 2 (SARS-CoV-2) a pandemic less than three months after its initial emergence in Wuhan, China [1,2]. SARS-CoV-2 is a zoonotic Betacoronavirus transmitted through person-person contact through airborne and fecal-oral routes, and has caused over 693,000 confirmed coronavirus disease 2019 (COVID-19) cases and 33,000 associated deaths worldwide [2–5]. While there is limited understanding of SARS-CoV-2 pathogenesis, extensive studies have been performed on how its closely related cousins, SARS-CoV and MERS-CoV (Middle East respiratory syndrome-related coronavirus), invade host cell. Upon initially contacting the surface of a host cell, SARS-CoV and MERS-CoV exploit host cell proteases to prime their surface spike glycoproteins (SGPs) for fusion activation, which is achieved by receptor binding, low pH, or both [6,7]. The receptor binding domain (RBD) resides within subunit 1 (S1) while subunit 2 (S2) facilitates viral-host cell membrane fusion [6]. Activated SGP undergoes a conformational change followed by an initiated fusion reaction with the host cell membrane [6]. Endocytosed virions are further processed by the endosomal protease cathepsin L in the late endosome [7,8]. Both MERS-Cov and SARS-CoV require proteolytic cleavage at their S2’ site, but not at their S1-S2 junction, for successful membrane fusion and host cell entry [6,7]. Additionally, receptors involved in fusion activation of SARS-CoV and MERS-CoV include heparan sulfate (HS) and angiotensin-converting enzyme 2 (ACE2), and dipeptidyl peptidase 4 (DPP4), respectively [9–11].

SARS-CoV and other pathogens arrive at a host cell surface by clinging, through their surface proteins, to linear, sulfated polysaccharides called glycosaminoglycans (GAGs) [12–14]. The repeating disaccharide units of GAGs, comprised of a hexosamine and a uronic acid or a galactose residue, are often sulfated (S1 Fig) [15]. GAGs are generally found covalently linked to core proteins as proteoglycans (PGs) and reside inside the cell, at the cell surface, and in the extracellular matrix (ECM) [15]. GAGs facilitate various biological processes, including cellular signaling, pathogenesis, and immunity, and possess diverse therapeutic applications [15]. For example, an FDA approved anticoagulant heparin (HP) is a secretory GAG released from granules of mast cells during infection [15,16]. Some GAG binding proteins can be identified by amino acid sequences known as Cardin-Weintraub motifs corresponding to ‘XBBXBX’ and ‘XBBBXXBX’, where X is a hydropathic residue and B is a basic residue, such as arginine and lysine, responsible for interacting with the sulfate groups present in GAGs [17,18]. Examination of the SARS-CoV-2 SGP sequence revealed that the GAG-binding motif resides within S1-S2 proteolytic cleavage motif (furin cleavage motif BBXBB) that is not present in SARS-CoV or MERS-CoV SGPs (Fig 1, S2 Fig, S3 Fig) [19]. Additionally, we discovered GAG-binding-like motifs within RBD and S2’ proteolytic cleavage site in SARS-CoV-2 SGP (Fig 1, S2 Fig, S3 Fig). This discovery prompted us to hypothesize that GAGs may contribute to SARS-CoV-2 fusion activation and host cell entry as a novel mechanism through SGP binding. We performed surface plasmon resonance (SPR)-based binding assays to determine binding kinetics of the interactions between various GAGs and SARS-CoV-2 SGP in comparison with SARS-CoV, and MERS-CoV SGP to address this question. Lastly, we performed blind docking on the trimeric SARS-CoV-2 SGP model to objectively identify the preferred binding GAG-binding sites on the SGP.

**Fig 1.**
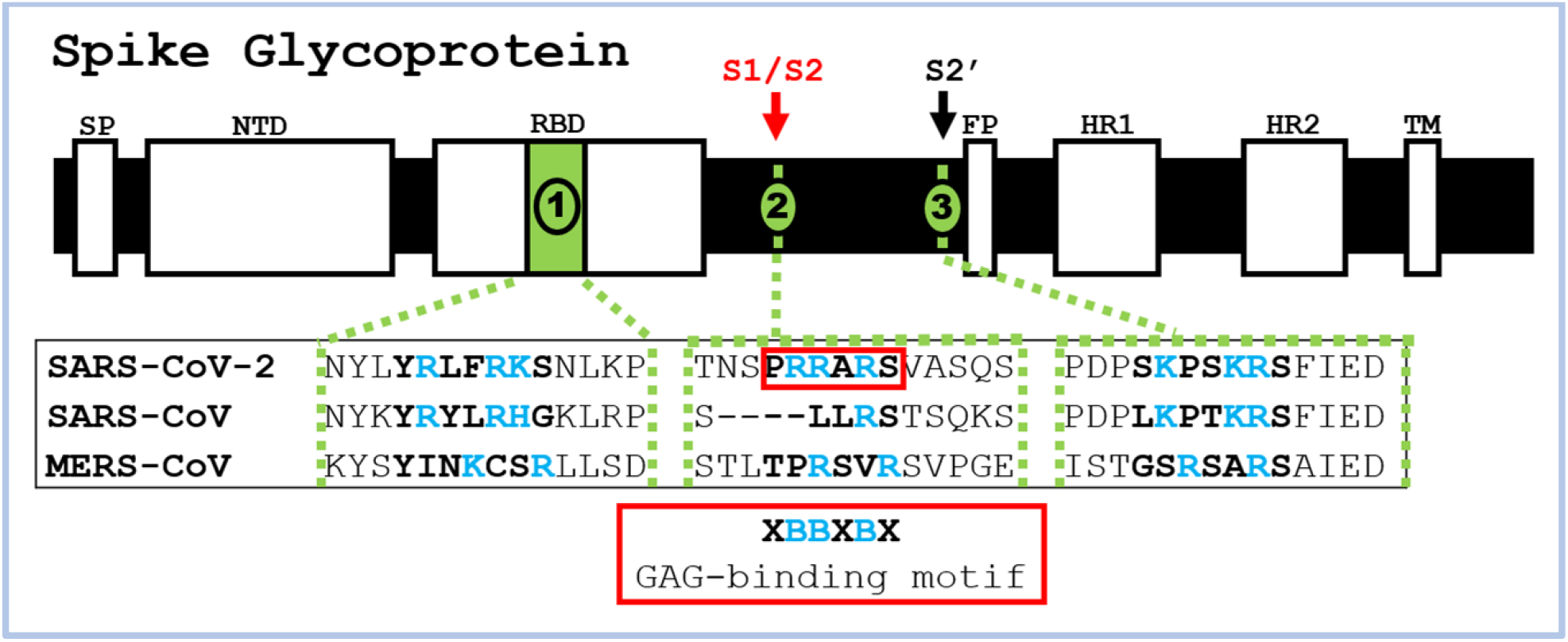
Identification of GAG-binding motif within SARS-CoV-2, SARS-CoV, and MERS-CoV SGPs. Domains in SGP include signal peptide (SP), N-terminal domain (NTD), receptor-binding domain (RBD), fusion peptide (FP), heptad repeat 1/2 (HR 1/2).

## Results

### Kinetic measurements of CoV SPG-HP interactions

Previous reports showed that various CoV bind GAGs through their SGPs to invade host cells [13]. In the current study, we utilized SPR to measure the binding kinetics and interaction affinity of monomeric and trimeric SARS-CoV-2, monomeric SARS-CoV and MERS-CoV with SGP-HP using a sensor chip with immobilized HP. Sensorgrams of CoV SGP-HP interactions are shown in Fig 2. The sensorgrams were fit globally to obtain association rate constant (*k_a_*), dissociation rate constant (*k_d_*) and equilibrium dissociation constant (K_D_) (Table 1) using the BiaEvaluation software and assuming a 1:1 Langmuir model. SARS-CoV-2 and MERS CoV SGP exhibited a markedly low dissociation rate constant (*k_d_ ~* 10^−7^ 1/s) suggesting excellent binding strength. The HP binding properties of monomeric SARS-CoV-2 SGP was comparable to that of the trimeric form (K_D_ of monomer and trimer were 40 pM and 73 pM, respectively). In comparison, previously known HP binding SARS-CoV SGP showed nearly 10-fold lower affinity, 500 nM. The extremely high binding affinity of SARS-CoV-2 SGP to HP was supported by the chip surface regeneration conditions. The immobilized HP surface could only be regenerated using a harsh regeneration reagent, 0.25% SDS, instead of the standard 2M NaCl solution used for removing HP-binding proteins. One reason for SARS-CoV-2 SGP monomer and trimer extremely high affinity to immobilized heparin is the high density of surface bound ligands might promote polyvalent interactions. The difference of binding kinetics and affinity of CoV SGPs to HP may also be due in part to the difference in protein sequence of the Cov SGPs. Based on amino acid alignment analysis using the Basic Local Alignment Search Tool (BLAST), SARS-CoV and SARS-CoV-2 SGPs share 76% similarity. Association rate constants (k_a_) for MERS-CoV SGP (339 (± 27) 1/M^−1^s^1^) was the lowest, followed by monomeric and trimeric SARS-CoV-2 SGP (2.5 × 10^3^ (± 62.7) M^−1^s^−1^ and 1.6 × 10^3^ (± 127) M^−1^s^−1^, respectively) (Table 1). SARS-CoV SGP had the highest K_a_, which was 4.12 × 10^4^ (± 136) M^−1^s^−1^. The differences in k_a_ values suggest a different mechanism when each SGP binds HP in addition to differences in binding strengths.

**Table 1.**
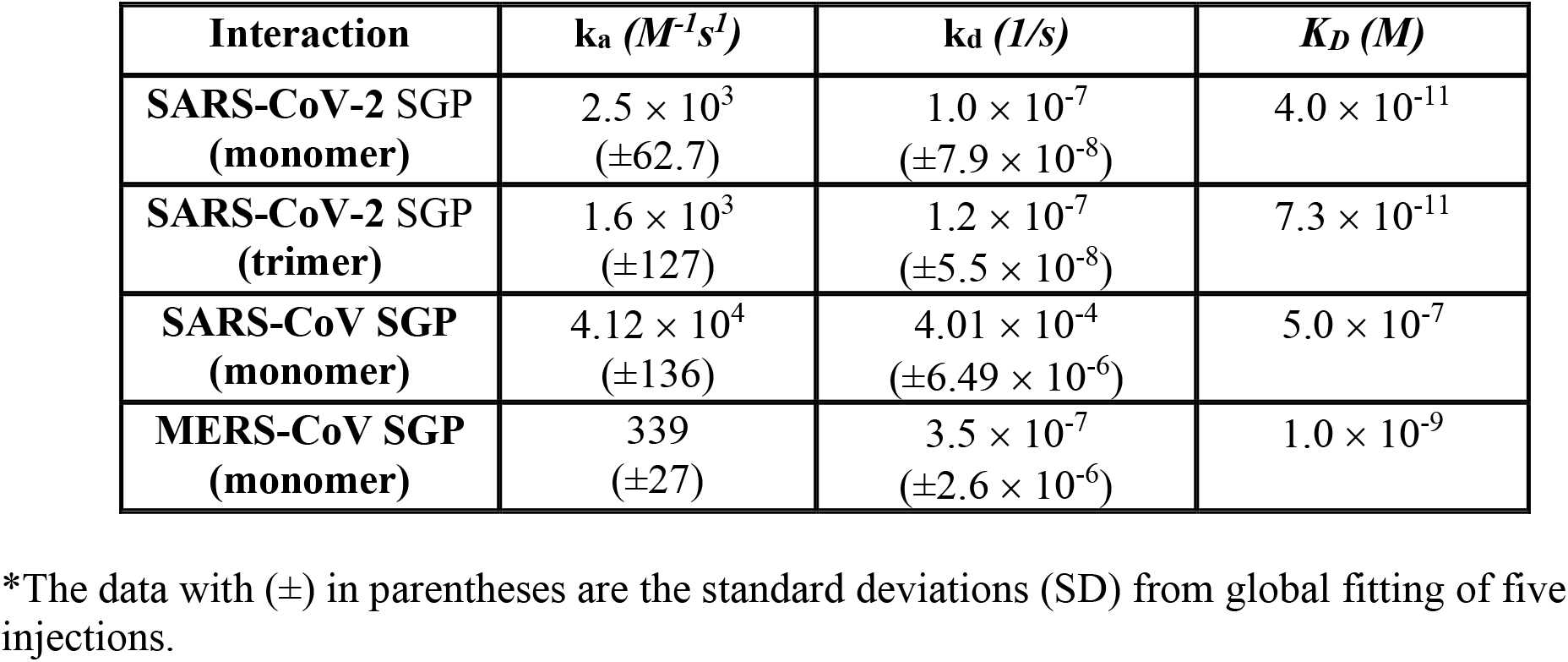
Summary of kinetic data of CoV SGP-HP interactions*.

**Fig 2.**
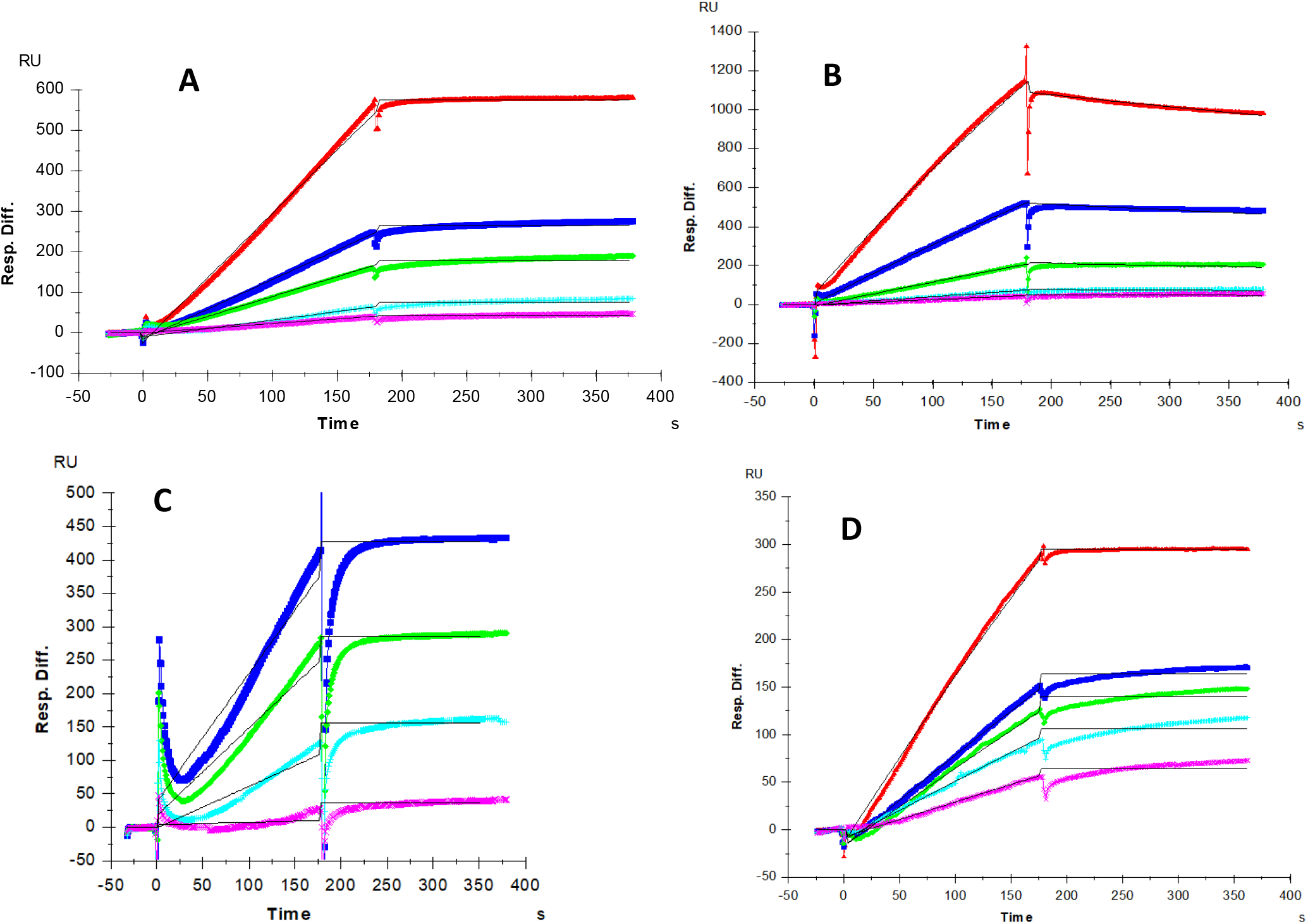
SPR sensorgrams for binding kinetics/affinity measurements for SGP-HP interactions. (A) SARS-CoV-2 SGP (monomer), concentration of SGP (from top to bottom): 100, 50, 25, 12.5 and 6.25 nM. (B) SARS-CoV SGP, concentrations of SARS-CoV SGP (from top to bottom): 100, 50, 25, 12.5 and 6.25 nM. (C) MERS CoV SGP, concentrations of MERS CoV SGP (from top to bottom): 100, 50, 25, 12.5 and 6.25 nM. (D) SARS-CoV-2 SGP (trimer), concentration of SGP (from top to bottom): 800, 400, 200, 100 and 50 nM. The black curves are the fits using a 1:1 Langmuir model from BIAevaluate 4.0.1.

### SPR solution competition study on the interaction between surface-bound HP and SARS-CoV-2 SGP to HP-derived oligosaccharides in solution

Solution/surface competition experiments were performed by SPR to examine the effect of the saccharide chain length of HP on the SARS-CoV-2 SGP-HP interaction. HP-derived oligosaccharides of different lengths, from tetrasaccharide (dp4) to octadecasaccharide (dp18), were used in these competition studies. The same concentration (1000 nM) of HP oligosaccharides were mixed in the SARS-CoV-2 SGP protein (50 nM)/ HP interaction solution. Negligible competition was observed (S4 Fig) when 1000 nM of oligosaccharides (from dp4 to dp18) were present in the protein solution suggesting that the SARS-CoV-2 SGP-HP interaction is chain-length dependent and it prefers to bind full chain (~dp30) HP.

### SPR solution competition study of different chemically modified HP derivatives and GAGs

Competition levels measured by SPR for chemically modified HP derivatives are shown in Fig 3. The results of these studies demonstrate that all the chemically modified HPs, *N*-desulfated HP, 2-*O*-desulfated HP and 6-*O*-desulfated HP, were unable to compete with immobilized HP for binding to SARS-CoV-2 SGP-HP suggesting all the sulfate groups within HP have critical impact on this interaction.

**Fig 3.**
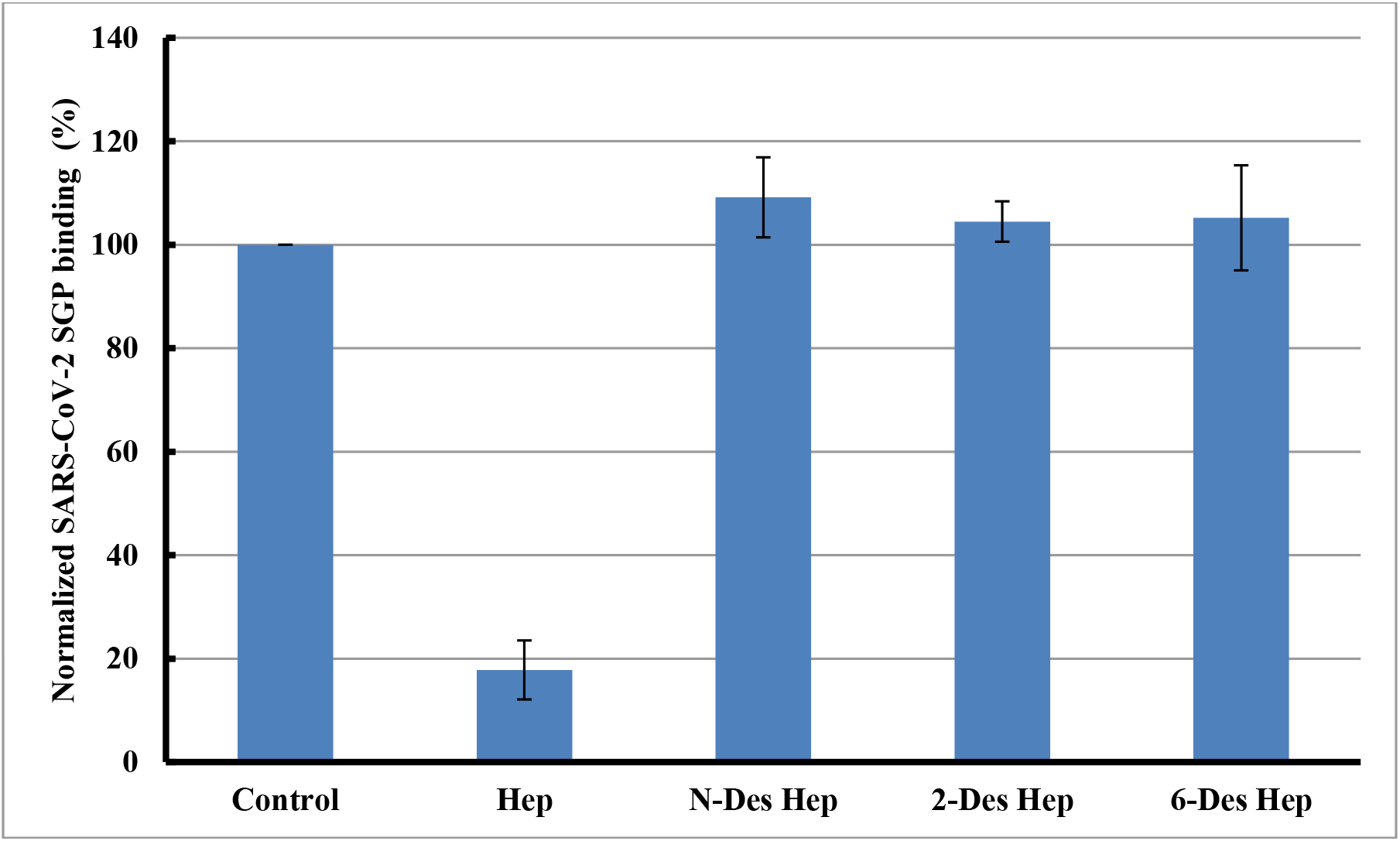
Bar graphs of normalized SARS-CoV-2 SGP binding preference to surface HP by competing with different chemical modified HP in solution. Concentration was 50 nM for SARS-CoV-2 SGP and 1000 nM for different chemical modified HP. All bar graphs based on triplicate experiments.

SPR competition assay was also used to test the binding preference of SARS-CoV-2 SGP for various GAGs (S1 Fig), including various chondroitin sulfates, dermatan and keratan sulfates, and the results are shown in S5 Fig. Weak or no inhibitory activities were observed for all GAGs tested, suggesting that the binding of SARS-CoV-2 SGP protein to GAGs appears to be HP specific and greatly influenced by the level of sulfation within the GAG.

### SPR solution competition dose response analysis of HP, Tri-sulfated HS and NACH

Solution competition dose response analysis between surface immobilized HP and various soluble glycans (HP, non-anticoagulant trisulfated (TriS) HS, and non-anticoagulant low molecular weight HP (NACH)) was performed to calculate their IC_50_ values (Figs 4A-4E). SARS-CoV-2 SGP protein (50 nM) samples were pre-mixed with different concentrations of glycans before injection into the HP chip. The sensorgrams (Figs 4A, 4C and 4E) show that once the active binding sites on SARS-CoV-2 SGP were occupied by glycans in solution, the binding of SARS-CoV-2 SGP to the surface-immobilized HP decreased resulting in a reduction of signal in a concentration dependent fashion. The IC_50_ values (concentration of competing analyte resulting in a 50% decrease in RU) were calculated from the plots SARS-CoV-2 SGP binding signal (normalized) vs. glycans concentration in solution (Figs 4B, 4D and 4F). The IC_50_ values of HP, TriS HS and NACH were 0.056 μM, 012 μM and 26.4 μM, respectively.

**Fig 4.**
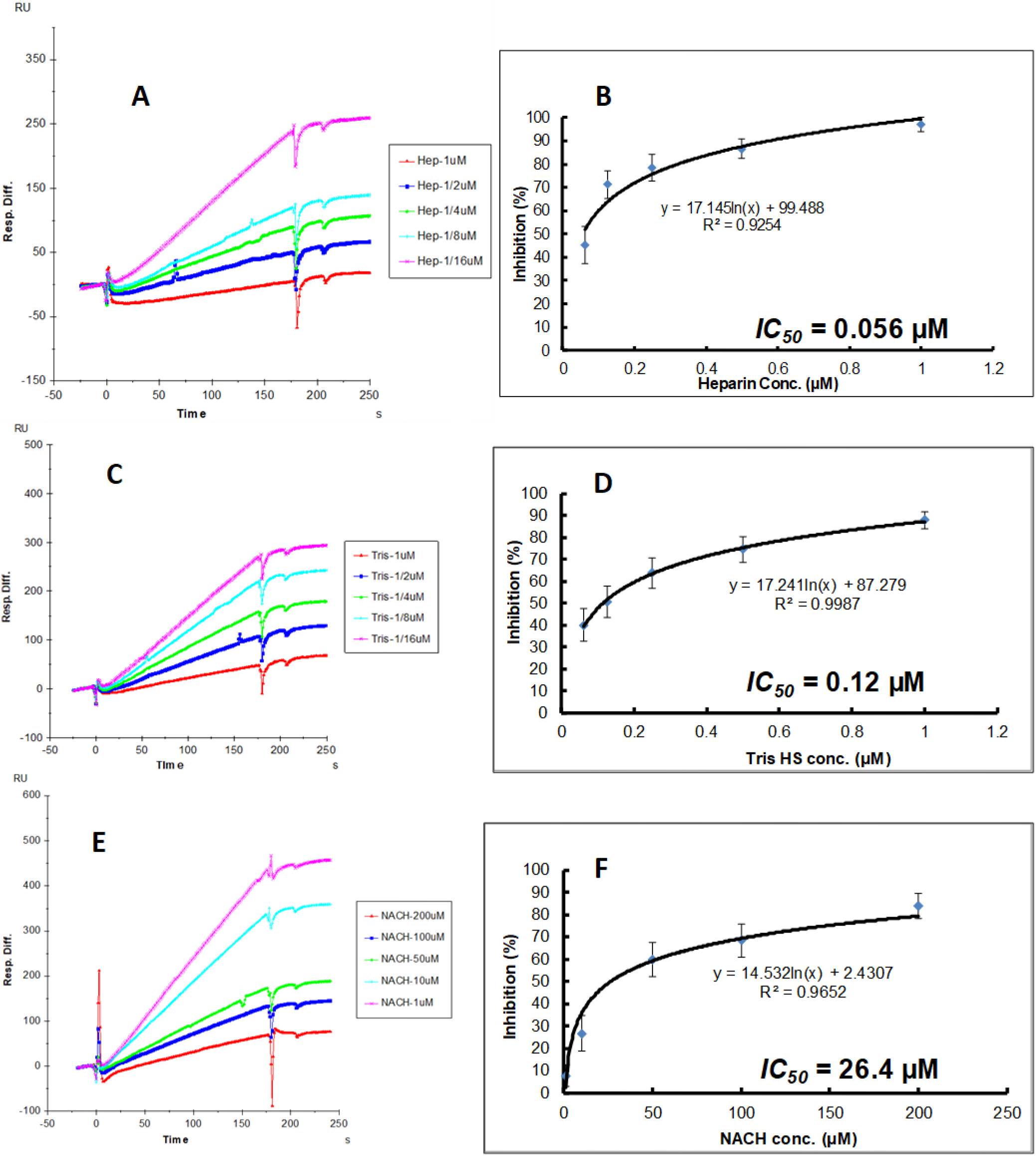
Inhibition analysis of glycans on the interactions between SARS-CoV-2 SGP and HP using SPR; SARS-CoV-2 SGP concentration was 50 nM. (A) Competition SPR sensorgrams of SARS-CoV-2 SGP-HP interaction inhibiting by different concentration of heparin. (B) Dose response curves for IC_50_ calculation of heparin using SARS-CoV-2 SGP inhibition data from surface competition SPR. (C) Competition SPR sensorgrams of SARS-CoV-2 SGP-HP interaction inhibiting by different concentration of TriS HS. (D) Dose response curves for IC_50_ calculation of TriS HS using SARS-CoV-2 SGP inhibition data from surface competition SPR. (E) Competition SPR sensorgrams of SARS-CoV-2 SGP-HP interaction inhibiting by different concentration of NACH. (F) Dose response curves for IC_50_ calculation of NACH using SARS-CoV-2 SGP inhibition data from surface competition SPR.

### Identification of GAG-binding motifs by blinding docking analysis

Using a modified version of Autodock Vina tuned for use with carbohydrates (Vina-Carb) [20,21], we performed blind docking on the trimeric SARS-CoV-2 SGP model to discover objectively the preferred binding GAG-binding sites on the SGP protein surface. The SGP contains three putative GAG-binding motifs with the following sequences: 453-459 (YRLFRKS), 681-686 (PRRARS), and 810-816 (SKPSKRS), which we define as sites 1, 2, and 3, respectively (Fig 1, S2 Fig, S3 Fig). An HS hexasaccharide fragment (GlcA(2S)-GlcNS(6S)) binds site 2 in each monomer chain in the trimeric SGP (Fig 5C, S3 Fig). The docking results also indicates that HS may bind to site 1 when the apex of the S1 monomer is in an open conformation, as this allows basic residues to be more accessible to ligand binding. The site 1 residues are less accessible for GAG binding when the domain is in a closed conformation (Fig 5D). The electrostatic potential surface representation of the trimeric SGP confirms that the GAG-binding poses generally prefer regions of positive charge, as expected, and illustrates that basic residues within site 3 are not exposed for binding to HS on any of the chains (Fig 5A). Finally, our blind docking analysis reveals that a longer HS polymer may span an inter-domain channel that contains site 2.

**Fig 5.**
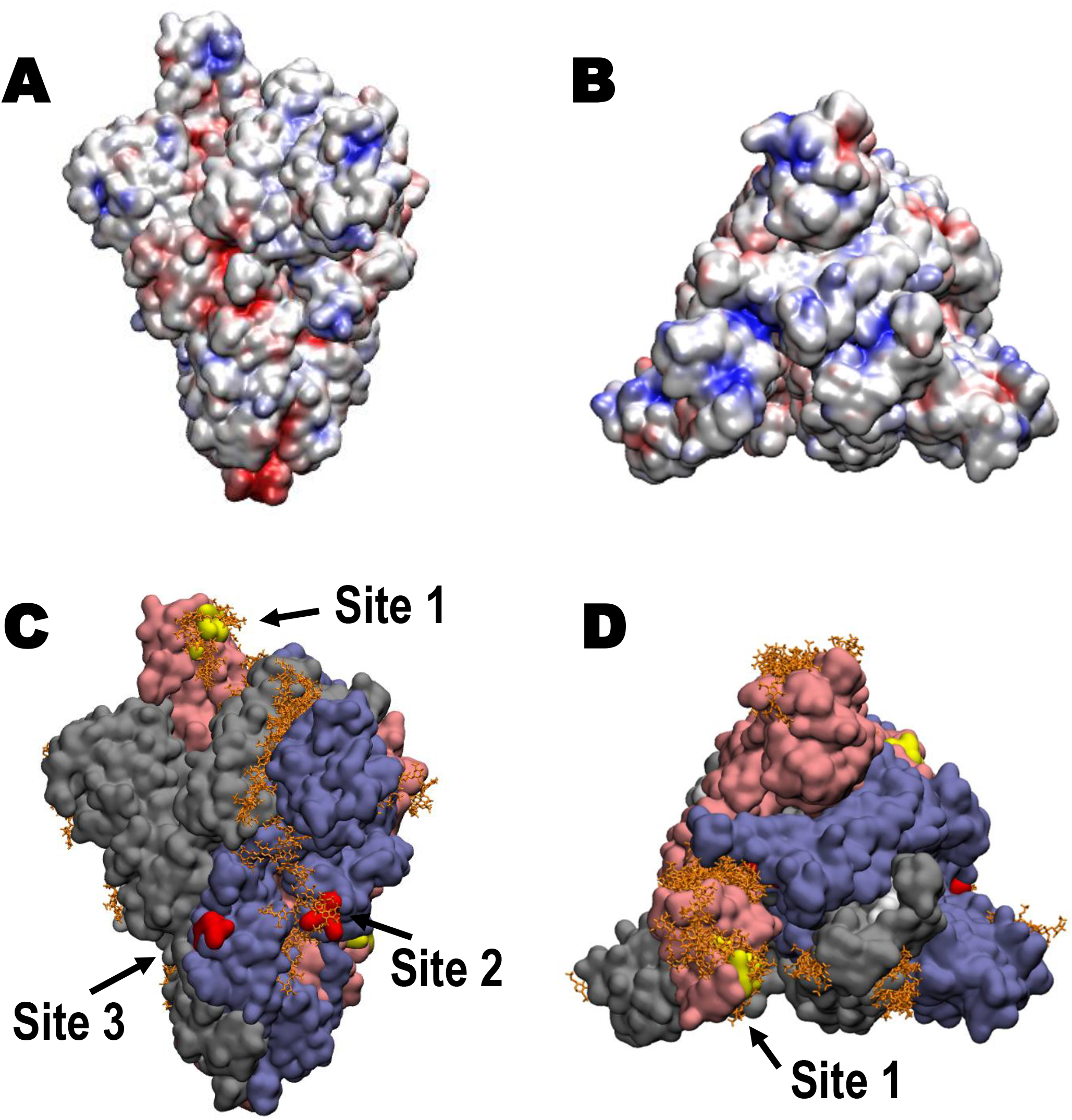
Structure of trimeric SARS-CoV-2 SGP and proposed GAG-binding motifs. (A) Electrostatic potential surface (-ve charge (red) to +ve charge (blue)) computed with Chimera. (B) Electrostatic potential surface showing a top view of the SGP trimer. (C) Solvent accessible surface of the SARS-CoV-2 SGP trimer (pink (Chain A), grey (Chain B), blue (Chain C)) showing the predicted poses of HS hexasaccharides (orange) obtained from unbiased docking, and the three GAG-binding motifs (yellow (Chain A), white (Chain B), red (Chain C)), image generated with VMD [37]. (D) Solvent accessible surface showing a top view of the SGP trimer. Amino acid sequences for GAG-binding motifs site 1, 2, and 3 are YRLFRKS, PRRARS, and SKPSKRS.

## Discussion

The original SARS-CoV and numerous pathogens exploit host cell surface GAGs during the initial step of host cell entry [12–14]. Based on our discovery of GAG-binding and GAG-binding-like motifs at site 1 (within the RBD, Y453-S459), site 2 (at the proteolytic cleavage site at S1/S2 junction, P681-S686), and site 3 (at the S2’ proteolytic cleavage site, S810-S816), we hypothesized that SARS-CoV-2 may also interact with host cell surface GAGs through its SGPs to invade host cell (Fig 1, S2 Fig, S3 Fig). The predominant GAG in normal human lung is HS followed by CS [22] and it is noteworthy that lung tissue is rich in mast cells and has been a source of commercial HP [23]. Using unbiased docking, we found that TriS HS hexasaccharide (GlcA(2S)-GlcNS(6S)) binds site 2 in each monomer chain in the trimeric SARS-CoV-2 SGP (Fig 5C). TriS HS hexasaccharide may additionally bind site 1 when the SGP monomer is in an open conformation, but not site 3 where the basic residues are not accessible to the surface (Figs 5C and 5D). Docking results indicated that the HS hexasaccharides could span an inter-domain channel that includes site 2, suggesting a mechanism for the binding of a longer HS sequence (Fig 5C).

Next, we experimentally determined binding kinetics for the interactions between HP (rich (60-80%) in TriS domains) and monomeric SARS-CoV-2, trimeric SARS-CoV-2, monomeric SARS-CoV, and monomeric MERS-CoV SGPs using SPR binding assays (Fig 2 and Table 1). GAG-protein interactions are mainly electrostatically driven [24], thus, HS-binding proteins generally bind HP due to its higher degree of sulfation [15]. We discovered that HP binds both monomeric and trimeric SARS-CoV-2 SGP with remarkable affinity (K_D_ = 40 pM and 73 pM, respectively) (Fig 2 and Table 1). This was unexpectedly tight binding for a GAG-protein interaction as even one of one of the prototypical HP-binding proteins, fibroblast growth factor 2 (FGF2), has a K_D_ of 39 nM [25]. In comparison, SARS-Cov and MERS-CoV SGPs also bind HP, however, much more weakly with binding strengths of K_D_ = 500 nM and 1 nM, respectively (Fig 2 and Table 1). While HS facilitates SARS-CoV host cell entry and is an essential host cell surface receptor, its involvement in MERS-CoV host cell entry or binding kinetics for SARS-Cov and MERS-CoV SGPs had not previously been reported [10].

After discovering the high binding affinity between HP and SARS-CoV-2 SGP, we next found that the degree and position of sulfation within HP was important for its successful binding to monomeric SARS-CoV-2 SGP (Figs 3 and 4). *N*-, 2-*O*, and 6-*O*-sulfation were all required for binding to SARS-CoV-2 SGP (Fig 3). This was additionally demonstrated when competitive binding studies gave IC_50_ values of HP (0.056 μM), TriS HS (NS2S6S) (0.12 μM), and NACH (26.4 μM) for the inhibition of SARS-CoV-2 SGP binding to immobilized HP (Fig 4). Both HP and TriS HS are sulfated at *N*-, 2-*O*-, and 6-*O*-positions and have approximately the same molecular weight (and chain length) but HP has additional 3-*O*-sulfation, responsible for its anticoagulant activity (S1 Fig). NACH lacks an intact antithrombin binding site, has a lower average molecular weight of 5 kDa than HP, and a higher content (>90%) of TriS [26].

The low IC_50_ of these GAGs suggest that the FDA approved anticoagulant HP, or its non-anticoagulant derivatives, might have therapeutic potential against SARS-CoV-2 infection as competitive inhibitors. The location of proposed GAG-binding sites is also of interest. Unlike SARS-CoV and MERS-CoV SGPs, SARS-CoV-2 SGP has a novel insert in the amino acid sequence (681-686 (PRRARS)) that fully follows GAG-binding Cardin-Weintraub motif (XBBXBX) and a furin-cleavage motif (BBXBB) at the S1/S2 junction (Fig 1). This site was also shown to be a preferred GAG-binding motif by our unbiased docking study (Fig 5). Proteolytic cleavage at S1/S2 is not required for successful viral-host cellular membrane fusion in SARS-CoV and MERS-CoV SGPs [6,7]. Proteolytic cleavage primes the SGP for fusion activation and may additionally influence cell-cell fusion, host cell entry, and/or the infectivity of the virus [13,27].

Some CoVs, including mouse hepatitis virus (MHV) and infectious bronchitis virus (IBV), possess both GAG-binding and furin cleavage motifs at their S1/S2 junction in their SGPs [13]. In the cases of MHV and IBV spike proteins, a single amino acid mutation near the GAG-binding and furin cleavage motifs resulted from a cell culture adaptation, and determines whether a virion binds GAGs or exploits host cell surface protease, but not both [28]. While not within the CoV family, human immunodeficiency virus type 1 (HIV-1) requires HS-binding to achieve optimal furin processing because HS binding allows selective exposure of furin cleavage site [29]. While the idea of repurposing HP as COVID-19 therapeutic sounds appealing, further questions, including *in vitro* relevance of GAGs as host cell surface receptors, proteolytic processing of SGPs at S1/S2 junction, and their relationship in host cell entry and infectivity, must first be carefully evaluated.

Based on our findings, we propose a model on how GAGs may facilitate host cell entry of SARS-CoV-2 (Fig 6). First, virions land on the epithelial surface in the airway by binding to HS through their SGPs (Fig 6A). Host cell surface proteoglycans utilize their long HS chains to securely wrap around the trimeric SGP (Fig 6A). During this step, heavily sulfated HS chains span inter-domain channel containing GAG-binding site 2 on each monomer in the trimeric SGP and binds site 1 within the RBD in an open conformation (Fig 5). Host cell surface and extracellular proteases, such as furin and transmembrane serine protease 2 (TMPRSS2), may process site 2 (S1/S2 junction) and/or 3 (S2’) and GAG chains come off from site 2 upon cleavage (Fig 6B). HS and ACE2 binding to more readily accessible RBD containing site 1 may drive conformational change of SGP and activate viral-cellular membrane fusion [30]. Finally, SGP on the endocytosed virion may utilize an endosomal host cell protease, such as cathepsin L, to further execute viral-cellular membrane fusion. (Fig 6C).

**Fig 6.**
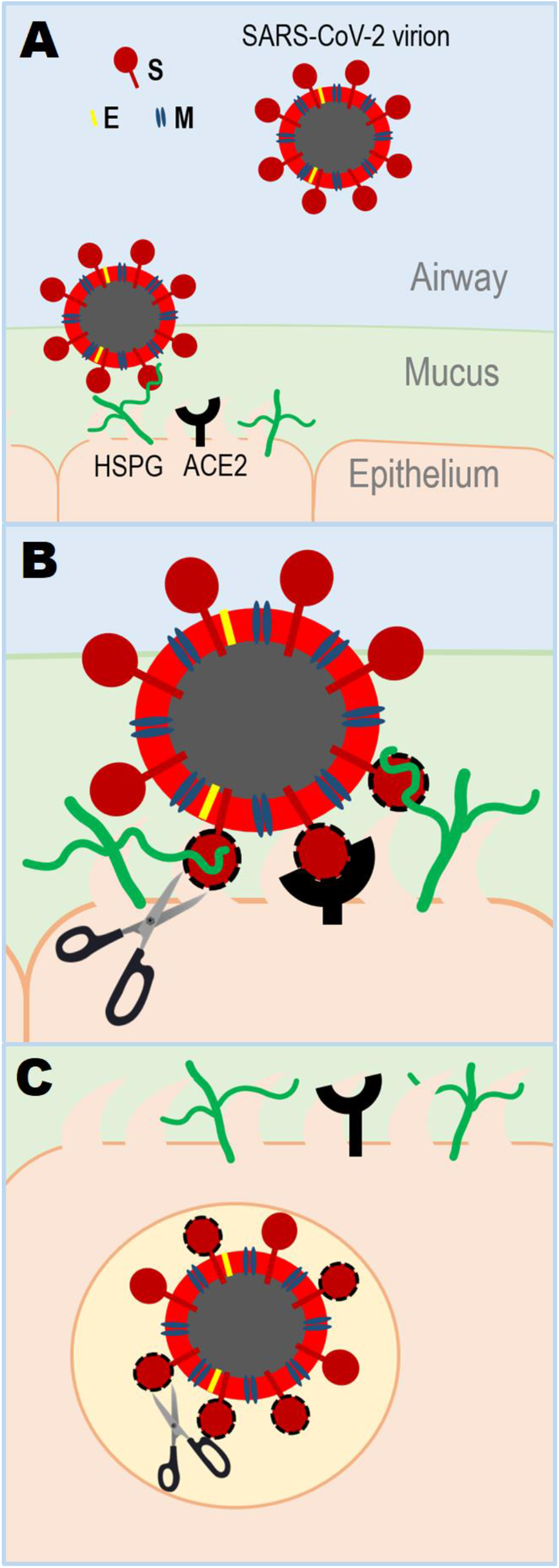
Proposed model of SARS-CoV-2 host cell entry. SARS-CoV-2 surface is decorated with envelop (E), membrane (M), and SGP [38]. (A) Virion lands on host cell surface by binding to heparan sulfate proteoglycan (HSPG). (B) SGP goes through proteolytic digestion by host cell surface protease, which initiates viral-host cell membrane fusion by conformational change caused by host cell receptor binding (HSPG and ACE2). (C) Virion enters the host cell and may further experience proteolytic processing by endosomal host cell protease.

In conclusion, we have discovered that GAGs can facilitate host cell entry of SARS-CoV-2 by binding to SGP in the current work. SPR studies demonstrate that both monomeric and trimeric SARS-CoV-2 SGP bind HP with remarkably high affinity and it prefers long, heavily sulfated (TriS rich) structures. Additionally, we reported low IC_50_ of HP and derivatives against HP and SARS-CoV-2 SGP interactions suggesting therapeutic potential of HP as COVID-19 competitive inhibitors. Lastly, unbiased computational ligand docking indicated that a TriS HS oligosaccharide preferably interacts with GAG-binding motifs at the S1/S2 junction and within receptor binding domain and hinted at mechanism of binding. This study adds to our current understanding of SARS-CoV-2 pathogenesis and serves a foundation for designing glycoconjugate vaccines and therapeutics to successfully contain and eliminate COVID-19.

## Materials and methods

### Materials

Monomeric SARS-CoV-2, SARS-CoV, and MERS-CoV SGPs were purchased from Sino Biological Inc. Trimeric SARS-CoV-2 SGP was kindly provided by Prof. Jason McLellan (University of Texas at Austin). The GAGs used in this study were porcine intestinal HP (HP) (average molecular weight, Mw = 16 kDa) and porcine intestinal heparan sulfate (HS) (Mw = 14 kDa) from Celsus Laboratories (Cincinnati, OH); chondroitin sulfate A (CSA, Mw = 20 kDa) from porcine rib cartilage (Sigma, St. Louis, MO), dermatan sulfate (DS, Mw = 30 kDa) from porcine intestine (Sigma), chondroitin sulfate C (CSC, Mw = 20 kDa) from shark cartilage (Sigma), chondroitin sulfate D (CSD, Mw = 20 kDa) from whale cartilage (Seikagaku, Tokyo, Japan) and chondroitin sulfate E (CSE, Mw = 20 kDa) from squid cartilage (Seikagaku). *N*-desulfated HP (*N*-DeS HP, Mw = 14 kDa) and 2-*O*-desulfated HP (2-DeS HP, MW = 13 kDa) were prepared in house based on protocols by Yates *et al* [31]. The 6-*O*-desulfated HP derivative, 6-DeS HP, Mw = 13 kDa, was generously provided by Prof. Lianchun Wang (University of South Florida). Non-anticoagulant low molecular weight HP (NACH) was synthesized from dalteparin, a nitrous acid depolymerization product of porcine intestinal HP, followed by periodate oxidation as described in our previous work [26]. TriS HS (NS2S6S) was synthesized from *N*-sulfo heparosan with subsequent modification with C5-epimerase and 2-*O*-and 6-*O*-sulfotransferases (2OST and 6OST1/6OST3) [32]. HP oligosaccharides included tetrasaccharide (dp4), hexasaccharide (dp6), octasaccharide (dp8), decasaccharide (dp10), dodecasaccharide (dp12), tetradecasaccharide (dp14), hexadecasaccharide (dp16) and octadecasaccharide (dp18) and were prepared from porcine intestinal HP controlled partial heparin lyase 1 treatment followed by size fractionation. The chemical structures of the GAGs are shown in S1 Fig. Sensor SA chips were from GE Healthcare (Uppsala, Sweden). SPR measurements were performed on a BIAcore 3000 operated using BIAcore 3000 control and BIAevaluation software (version 4.0.1).

### Preparation of HP biochip

Biotinylated HP was prepared by conjugating its reducing end to amine-PEG3-Biotin (Pierce, Rockford, IL). In brief, HP (2 mg) and amine-PEG3-Biotin (2 mg, Pierce, Rockford, IL) were dissolved in 200 μl H_2_O, 10 mg NaCNBH_3_ was added. The reaction mixture was heated at 70 °C for 24 h, after that a further 10 mg NaCNBH_3_ was added and the reaction was heated at 70 °C for another 24 h. After cooling to room temperature, the mixture was desalted with the spin column (3,000 MWCO). Biotinylated HP was collected, freeze-dried and used for SA chip preparation. The biotinylated HP was immobilized to streptavidin (SA) chip based on the manufacturer’s protocol. The successful immobilization of HP was confirmed by the observation of a 200-resonance unit (RU) increase on the sensor chip. The control flow cell (FC1) was prepared by 2 min injection with saturated biotin.

### Measurement of interaction between HP and CoV SGP using BIAcore

SGP samples were diluted in buffer (0.01 M HEPES, 0.15 M NaCl, 3 mM EDTA, 0.005% surfactant P20, pH 7.4,). Different dilutions of protein samples were injected at a flow rate of 30 μL/min. At the end of the sample injection, the same buffer was flowed over the sensor surface to facilitate dissociation. After a 3 min dissociation time, the sensor surface was regenerated by injecting with 30 μL of 0.25% sodium dodecyl sulfonate (SDS) to get fully regenerated surface. The response was monitored as a function of time (sensorgram) at 25 °C.

### Solution competition study between HP on chip surface and HP, HP-derived oligosaccharides, chemically modified HP or GAGs in solution using SPR

SARS-CoV-2 SGP (50 nM) mixed with 1 μM HP, HP-derived oligosaccharides, chemically modified HP or GAGs in SPR buffer were injected over HP chip at a flow rate of 30 μL/min, respectively. After each run, the dissociation and the regeneration were performed as described above.

### SPR solution competition IC_50_ measurement of glycans (HP, TriS HS and NACH) inhibition on SARS-CoV-2 SGP-HP interaction

Solution competition studies between surface HP and soluble glycans (HP, TriS HS and NACH) to measure IC_50_ were performed using SPR [33]. In brief, SARS-CoV-2 S-protein (50 nM) samples alone or mixed with different concentrations of glycans in SPR buffer were injected over the HP chip at a flow rate of 30 μl/min, respectively. After each run, dissociation and regeneration were performed as described above. For each set of competition experiments, a control experiment (only protein without glycan) was performed to ensure the surface was completely regenerated.

### Protein modeling

The 3D coordinates for the SGP trimer (NCBI reference sequence YP_009724390.1) were downloaded from the SWISS-MODEL homology modeling server [34]. The selected model was generated with the Cryo-EM structure PDB ID 6VSB as a template, which has a 99.26% sequence identity and 95% coverage for amino acids 27 to 1146. The template and resulting model is the “prefusion” structure with one of the three receptor binding domains (Chain A) in the “up” or “open” conformation [30]. Cryo-EM studies have revealed that the SARS-CoV-2 SGP trimer exists in two conformational states in approximately equal abundance [35]. In one state, all SGP monomers have their hACE2-binding domain closed, and in the other, one monomer has its hACE2-binding domain open, where it is positioned away from the interior of the protein.

### Ligand docking

Initial coordinates for a hexasaccharide fragment of HS (GlcA(2S)-GlcNS(6S))_3_ were generated using the GAG-Builder tool [36] at GLYCAM-Web (glycam.org) and used for unbiased (blind) docking. A hexasaccharide was chosen as being sufficiently long to represent a typical GAG length found in protein co-complexes [36] and to avoid introducing so many degrees of internal flexibility that the efficiency of the docking conformational search algorithm was impaired. Docking was performed using a version of Vina-Carb [21] that has been modified to improve its performance for GAGs. A grid box with dimensions (x = 190, y = 223, z = 184 Å) was placed at the geometric center the protein enclosing its entire surface. Docking was performed with default values, with the following exceptions: exhaustiveness = 80, chi_cutoff = 2, and chi_coeff = 0.5. All sulfate and hydroxyl groups and glycosidic torsion angles were treated as flexible, resulting in 83 ligand poses.

## Acknowledgements

We appreciate Prof. Jason McLellan from University of Texas Austin for providing trimeric SARS-CoV-2 SGP. Additionally, we thank professor Lianchun Wang from University of South Florida for providing 6-*O*-desulfated HP derivative.

