## Supporting Information for "Glycosaminoglycan binding motif at S1/S2 proteolytic cleavage site on spike glycoprotein may facilitate novel coronavirus (SARS-CoV-2) host cell entry"

**S1 Fig. Chemical structure of various glycosaminoglycans and HP oligosaccharides.**

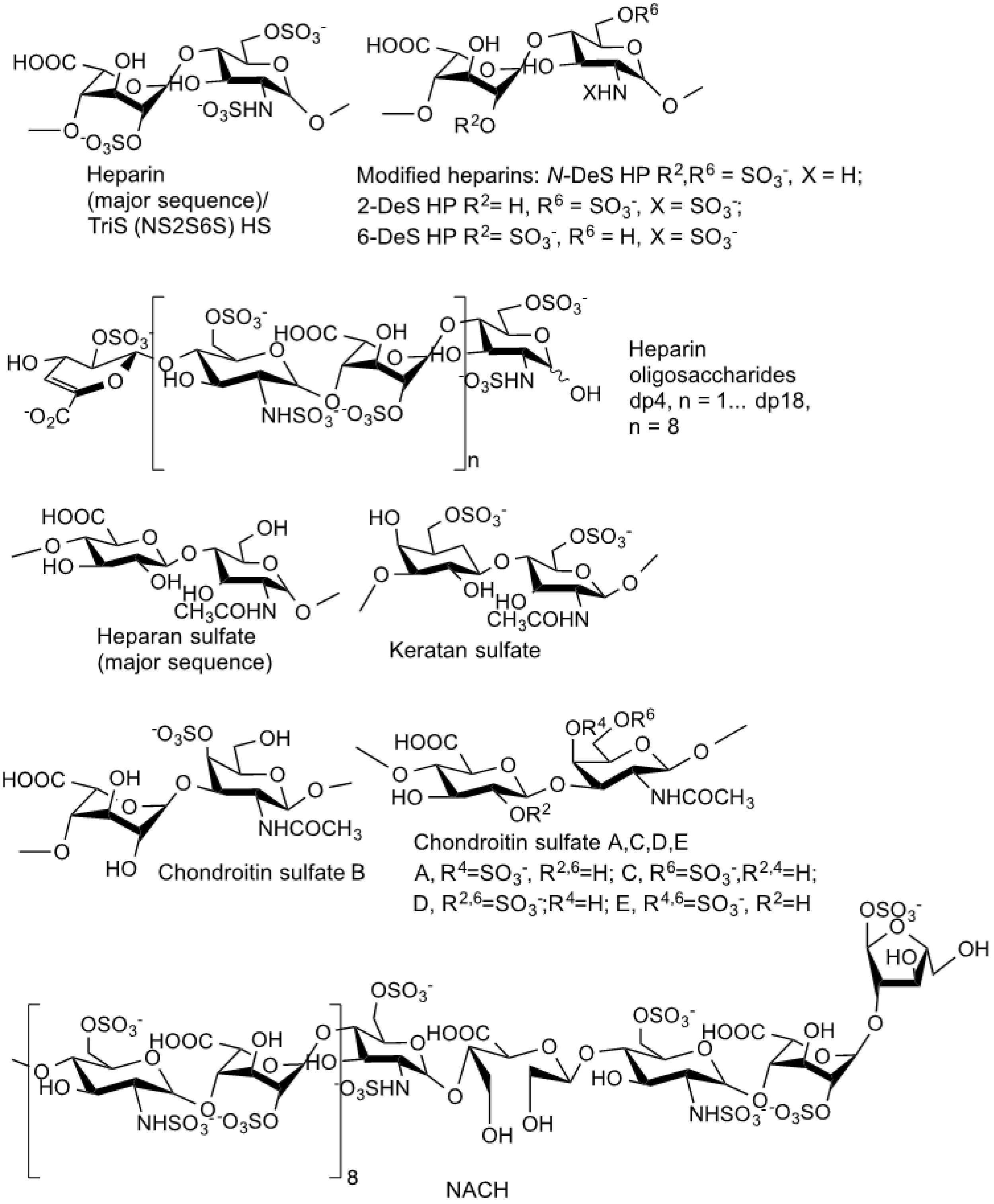

**S2 Fig. Multiple sequence alignment of SARS-CoV-2 (NC_045512.2), SARS-CoV (NC_004718.3), and MERS-CoV (NC_019843.3) SGP.** GAG-binding domains (GBD) and -like domains are boxed in red. Green regions represent receptor-binding domain (RBD). First and second arrows show proteolytic cleavage sites (PCS) at S1/S2 junction and S2' site, respectively. Fusion peptide (FP) is boxed in blue.

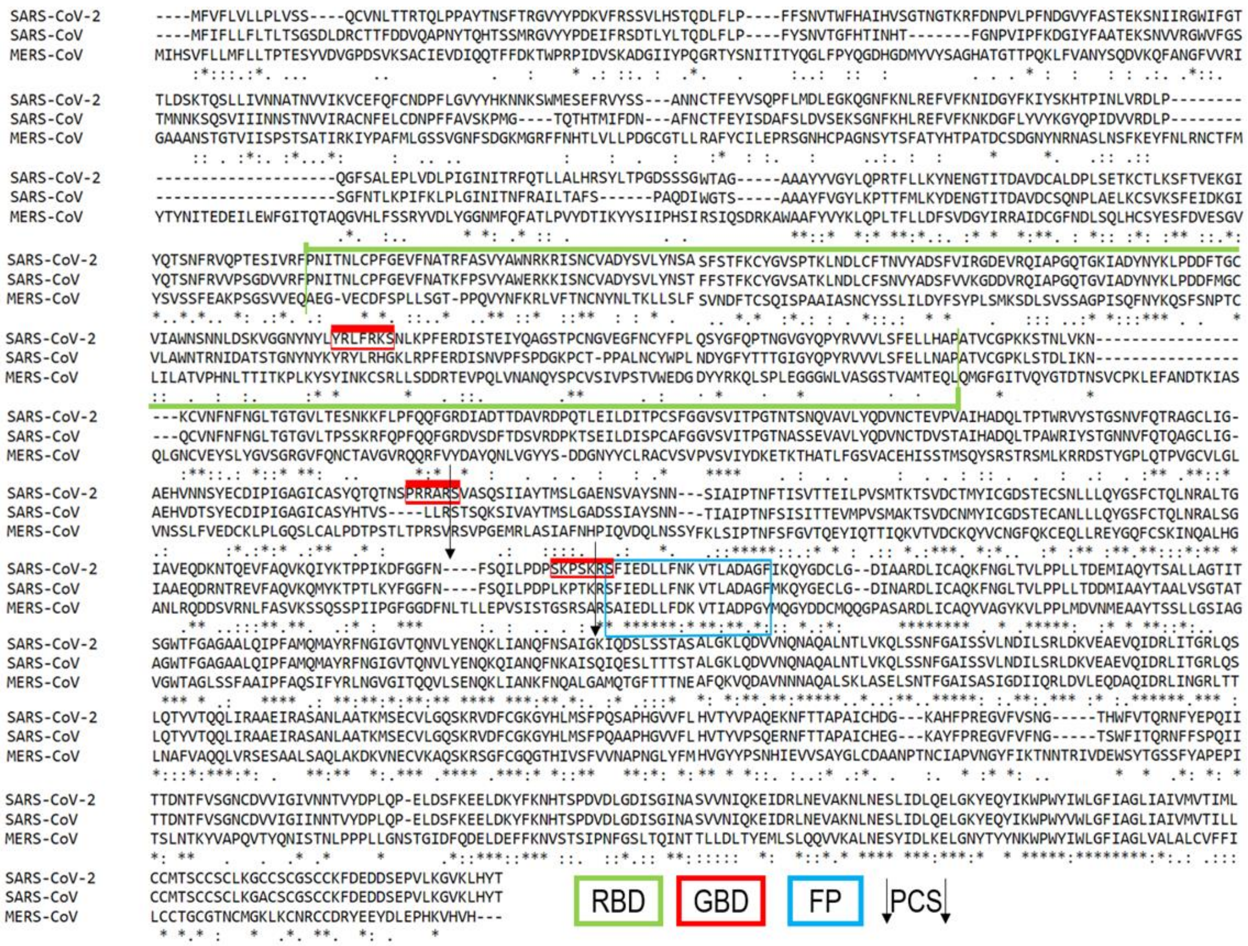

**S3 Fig. Structure of trimeric SARS-CoV-2 SGP and proposed GAG-binding motifs in solvent accessible surface ((A) and (C)) and in ribbon representations ((B) and (D)).** Chain A (pink), Chain B (grey), and Chain C (purple) are represented, and the atoms from the motif-based GAG binding sites are highlighted in yellow, white, and red, respectively. The GAG-binding motif on site 1 is accessible only when the monomer S1 domain adopts the "open" conformation (Chain A).

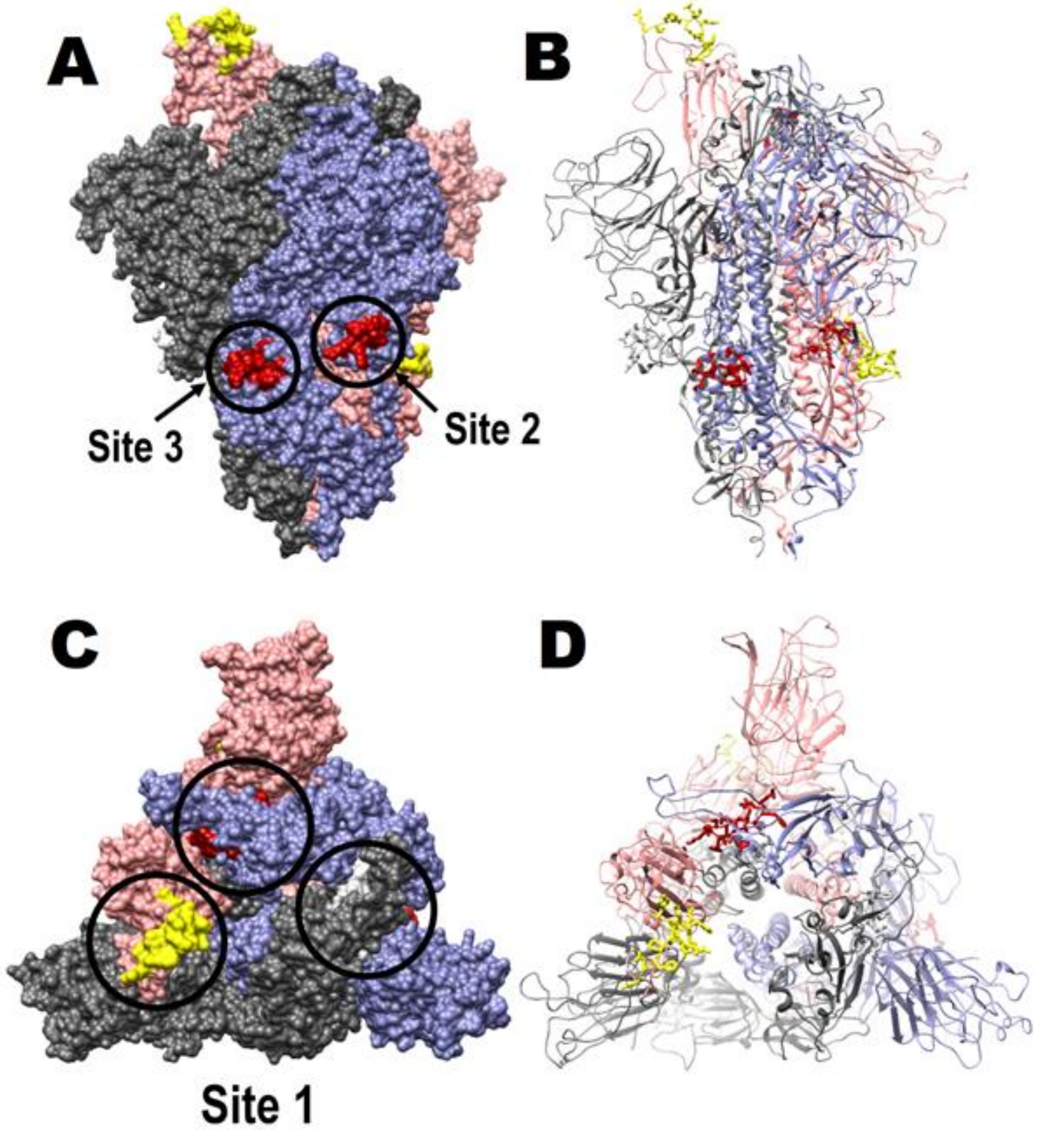

**S4 Fig. Bar graphs of normalized SARS-CoV-2 S-protein binding preference to surface heparin by competing with heparin-derived oligosaccharides in solution.** Concentration was 50 nM for SARS-CoV-2 S-protein and 1000 nM for heparin-derived oligosaccharides. All bar graphs based on triplicate experiments.

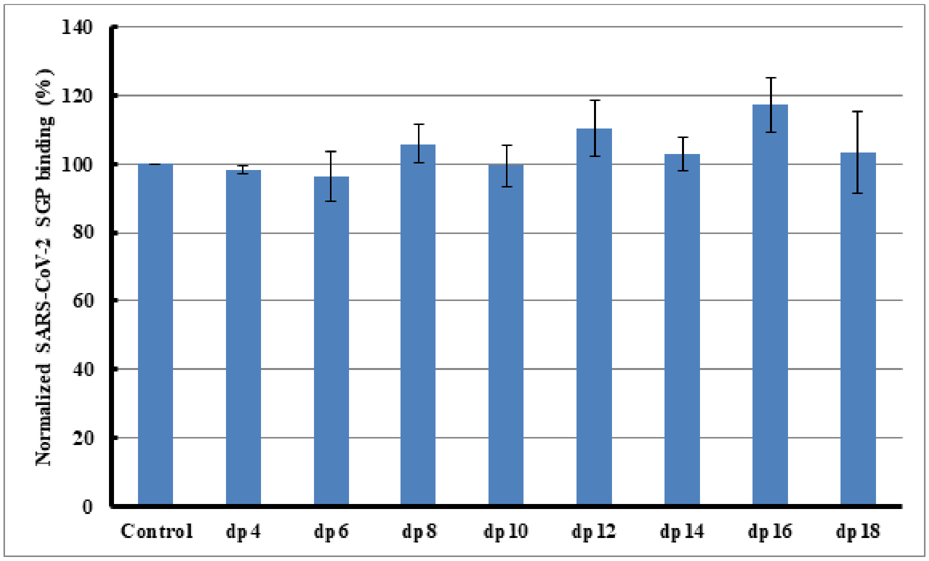

**S5 Fig. Bar graphs of normalized SARS-CoV-2 S-protein binding preference to surface heparin by competing with different GAGs in solution.** Concentration was 50 nM for SARS-CoV-2 S-protein and 1000 nM for different GAGs. All bar graphs based on triplicate experiments.

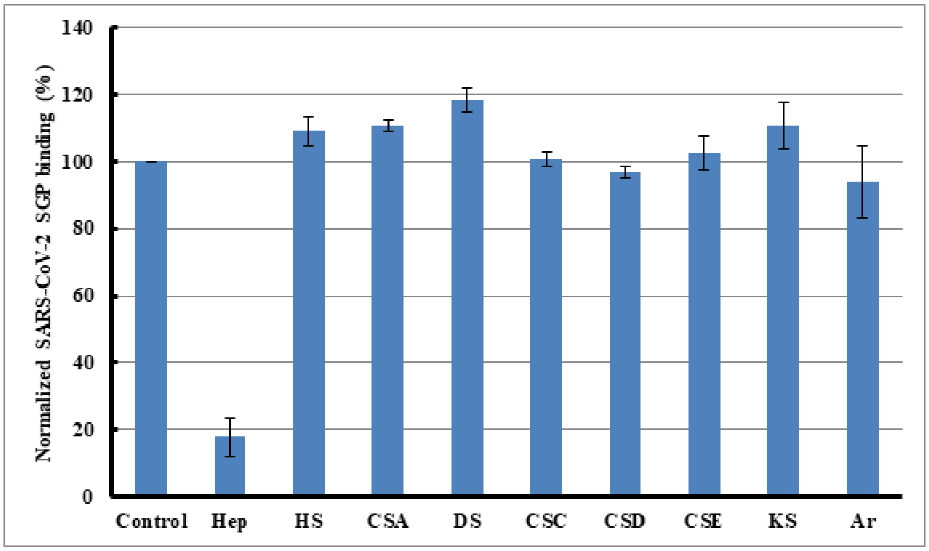
